# ATG8-interacting (ATI) 1 and 2 define a plant starvation-induced ER-phagy pathway and serve as MSBP1 (MAPR5) cargo-receptors

**DOI:** 10.1101/2020.01.29.924886

**Authors:** Jian Wu, Simon Michaeli, Gad Galili, Hadas Peled-Zehavi

**Author notes:** Correspondence: Hadas Peled-Zehavi Gad Galili.

## Abstract

ER-phagy, the selective autophagy of endoplasmic reticulum (ER) components, is known to operate in eukaryotes from yeast and unicellular algae to animals and plants. Thus far, only ER-stress derived ER-phagy was reported and analyzed in plants. In this study we characterize an ER-phagy pathway in *Arabidopsis thaliana* that is triggered by dark-induced starvation and not by ER-stress. This pathway is defined by the previously reported ATG8-interacting proteins, ATI1 and ATI2 and is regulated by the TOR signaling pathway. We further identified ER-localized Membrane Steroid Binding Protein 1 (MSBP1) as an ATI1 and 2 interacting protein and an autophagy cargo, and show that ATI1 and 2 serve as its cargo receptors. Together, these findings expand our knowledge on plants responses during energy deprivation and highlight the role of this special type of ER-phagy in this process.

## Introduction

Macroautophagy (hereafter referred to as autophagy) is a conserved eukaryotic catabolic mechanism for the removal and recycling of damaged or unneeded cellular components in response to developmental or environmental cues. It involves sequestration of cellular cargo in *de novo* formed double-membrane phagophores that finally close to form specialized vesicles called autophagosomes that are delivered to the lytic organelle, the vacuole in plants, for degradation [1, 2]. Unlike the ubiquitin-proteasome system, autophagy can handle large protein complexes and aggregates, as well as non-proteinous cargo such as nucleic acids, lipid bodies, and entire organelles [3]. The autophagy process requires the regulated and sequential involvement of a set of AuTophaGy-related (ATGs) proteins that are mostly conserved across eukaryotic organisms [3–6]. Autophagy can be a highly regulated selective turnover mechanism, targeting specific cellular components under specific cellular conditions. Recognition and recruitment of specific cargo to autophagosomes is mediated by cargo receptors. These receptors can recognize and bind the cargo directly or indirectly, and tether it to the forming autophagosome through interaction with core autophagy proteins [3, 7–9]. The ubiquitin-like ATG8 protein family plays an important role in selective autophagy. Soluble ATG8 is conjugated to the membrane lipid PE (phosphatidylethanolamine) on the expanding phagophore and is involved in autophagosome formation, trafficking and fusion with the vacuole. Additionally, ATG8 proteins function as hubs for selective autophagy routes, as they interact with multiple autophagy cargo receptors, mainly through an ATG8-interacting motif (AIM) or a newly identified ubiquitin-interacting motif (UIM) found on the receptors [3, 4, 10, 11]. In mammalian cells multiple cargo receptors have been identified in recent years, allowing for greater mechanistic understanding of selective autophagy processes and their impact on cellular functions and pathologies [8, 12–14]. However, plants lack orthologs to many of the mammalian cargo receptors, and although organelle-specific autophagy, as well as other types of selective autophagy were clearly demonstrated in plants, our mechanistic understanding of these processes and the selective cargo receptors that are required for them is still limited [3, 15]. Examples of plant selective autophagy receptors include Neighbor of BRCA 1 (NBR1), a functional hybrid of mammalian p62 and NBR1 autophagy receptors, which targets ubiquitylated protein aggregates in plant stress responses [16, 17] and virus particles [18], the proteasome subunit RPN10, which targets inactive ubiquitinated 26S proteasomes [19] and Tryptophan-rich Sensory Protein (TSPO) that can target plasma-membrane based aquaporins [20].

The endoplasmic reticulum (ER) is a complex membrane bound compartment in eukaryotic cells, and its varied cellular functions require rapid adjustment of its shape, size and protein and lipid content in response to changing cellular needs. The involvement of the ER in autophagic processes is multifaceted. The ER is a site for the nucleation of phagophores and a source for autophagosomal membranes [21, 22]. However, the ER is also the target of a selective type of autophagy, termed reticulophagy or ER-phagy, that is induced in yeast, mammalian cells and plants in response to ER-stress and starvation [22–26]. The ATG8-interacting ER proteins Atg39 and Atg40 were identified as the cargo receptors that mediate ER-phagy in yeast. Atg39 localizes to the perinuclear ER, while Atg40 is enriched in the cortical and cytoplasmic ER, suggesting that each protein is needed for the degradation of a different ER subdomain [25]. In mammals, six ER-resident autophagy receptors were identified to date: FAM134B [27], the long isoform of RTN3 (RTN3L) [28], CCPG1 (cell cycle progression 1)[29], SEC62 [30], AT3 (atlastin GTPase 3) [31] and TEX264 (testis expressed gene 264) [32, 33]. The different receptors seem to differ in their function and ER subdomain localization. FAM134B regulates starvation-induced turnover of ER sheets, while RTN3L and ATL3 are important for starvation-induced degradation of ER tubules [27, 28, 31]. CCPG1 and Sec62 are involved in ER stress induced ER-phagy [29, 30]. The newest addition to the cohort of mammalian ER-phagy receptors, TEX264, is ubiquitously expressed and was suggested to be a major contributor to starvation-induced ER-phagy [32, 33].

In *Arabidopsis thaliana*, treatment of seedlings with the ER-stress inducing compounds tunicamycin (TM) and dithiothreitol (DTT) induces ER-phagy and delivery of ER fragments to the vacuole [24]. Induction of autophagy was also seen in Chlamydomonas exposed to TM and DTT treatments [34]. The ER stress regulator Inositol-Requiring Enzyme-1b (IRE1b) is required for ER-phagy induction in *Arabidopsis*. However, the known downstream target of IRE1b, the transcription factor bZIP60, is not required [24]. Recently, IRE1 was shown to participate in IRE1-dependent decay of mRNAs (RIDD), in which mRNAs are degraded by IRE1 upon ER stress. As some of these mRNAs were shown to inhibit autophagy upon overexpression, it was suggested that IRE1b might regulate the degradation of mRNAs that interfere with autophagy induction [35]. However, the mechanism that allows for selective ER-phagy is still unclear, and no autophagy cargo receptors were identified in plants, though the *Arabidopsis* homolog of Sec62, AtSec62, was recently suggested to play a role in ER-phagy [36]. Thus, it is not known whether ER-stress induced ER-phagy is selective for specific ER subdomains or components.

The plant-specific trans-membranal ATI1 and ATI2 proteins (Atg8-Interacting Protein 1 and 2) were identified in a yeast two hybrid screen for *Arabidopsis* ATG8f-interacting proteins [37]. They were shown to localize to dark-induced ER and chloroplast-associated bodies (ATI-bodies) that are transported to the vacuole [37, 38]. Although ATI-bodies are distinct from autophagosomes, the delivery of the ATI1 proteins to the vacuole requires the autophagy machinery, suggesting that ATI-bodies eventually associate with autophagosomes [38]. ATI-bodies were shown to deliver chloroplast-targeted green fluorescent protein (GFP) to the vacuole and ATI1 can bind both stromal and membrane-bound chloroplast proteins [38]. Thus, it was suggested that the ATI proteins are chlorophagy cargo receptors [39]. However, their ER-related function is not clear. Recently, ATI1 and ATI2 were shown to be involved in an endogenous autophagic degradation pathway of ER-associated ARGONAUTE1 (AGO1) that is significantly induced following expression of the viral suppressor of RNA silencing P0 protein [40]. As ATI1 directly interacts with AGO1, it was suggested that it might function as selective cargo receptor for ER-localized AGO1. In this study we show that ATI1 and 2 define a novel type of dark-induced and TOR dependent ER-phagy pathway in *Arabidopsis.* We further show that ER-localized Membrane Steroid Binding Protein1 (MSBP1) interacts with the ATI proteins and is degraded by autophagy. Notably, ATI1 is required for the autophagic turnover of MSBP1, supporting the role of the ATI proteins as autophagy cargo receptors involved in selective ER-phagy.

## Results

### ATI1 is involved in ER-phagy in response to carbon starvation

To look at the possible involvement of the ATI proteins in ER-phagy, we used a transgenic line co-expressing an ER marker (mCherry-HDEL) and ATI1 fused to GFP (ATI1-GFP) [37]. Leaves of 4-5 week old plants were treated with the vATPase inhibitor concanamycin A (conA), that stabilizes autophagic bodies by raising the vacuolar pH, and the plants were either left under regular light conditions or darkened for 24 h. ATI1-labeled bodies or the ER marker were rarely observed in the vacuoles of leaves under regular light conditions (Figure 1A). However, in accordance with previous results [37, 38], numerous ATI1-labeled bodies were observed in the vacuole lumen after 24 h dark treatment (Figure 1B, left panel). Puncta labeled by the ER marker were also clearly visible in the vacuole following dark and conA treatment (Figure 1B, middle panel), suggesting that ER-phagy took place under these conditions. Interestingly, following dark treatment, the ER marker was highly co-localized with ATI1 in the vacuole (Figure 1B, right panel). Furthermore, the ER marker was surrounded by the membranal ATI1 that generated ring-like structures (Figure 1B, inset), suggesting that ATI1 is involved in ER-phagy in response to carbon starvation, and that ER components are delivered to the vacuole with ATI1. Similar results were observed in seedlings that were transferred from solid growth media to liquid media without sucrose in the presence of conA and darkened for 24 h (Figure S1A and B). To look at the possible involvement of ATI1 in ER-stress induced ER-phagy we took advantage of the relative stability of GFP. Vacuolar degradation of proteins tagged with GFP was shown to result in loss of the tagged protein and release of the relative stable free GFP [2, 4]. Thus, increase in the free GFP / total GFP ratio is indicative of increased autophagic turnover of the tagged protein. Indeed, increased free GFP release from ATI1-GFP was observed following dark treatment of leaves (Figure 1C and 1D). However, following incubation of leaves with the glycosylation inhibitor tunincamycin (TM), that induces ER stress [41, 42], no increase in free GFP / total GFP ratio was observed (Figure 1C and 1D), although the expression of several unfolded protein response (UPR) target genes was elevated as expected (Figure S1C and S1D). Similar results were observed following TM treatment of seedlings (Figure S1E). Taken together, our results suggest that ATI1 is involved in carbon starvation induced ER-phagy, but not in ER stress induced ER-phagy.

**Figure 1.**
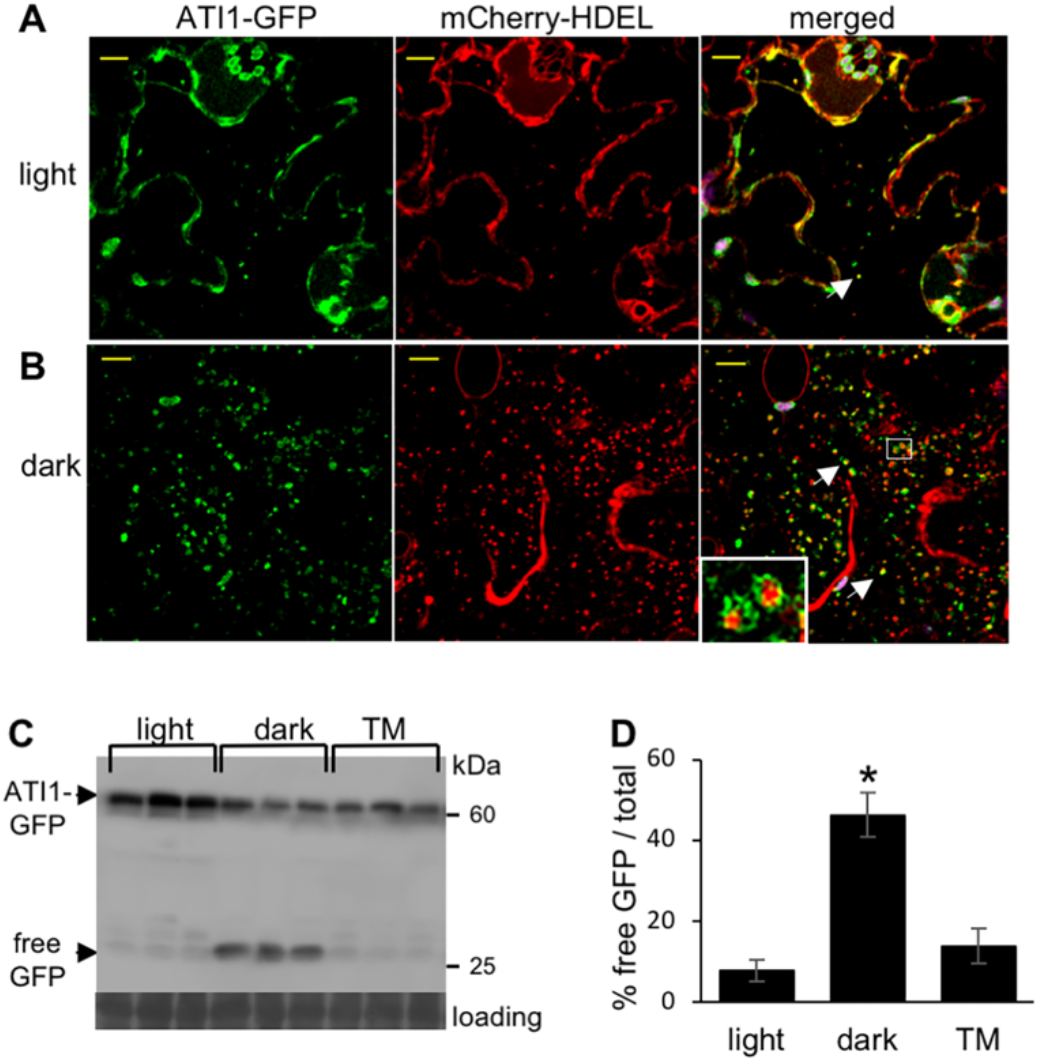
ATI1 is involved in ER-phagy in response to carbon starvation but not to ER stress. Leaves of ATI1-GFP/mCherry-HDEL co-expressing plants were infiltrated with conA and the plants were either left under regular light conditions **(A)** or darkened **(B)** for 24 h. Representative confocal images of leaves show ATI1-GFP in green, mCherry-HDEL in red and the overlay indicating co-localization in yellow (merged). Magnification of the area in the white rectangle is shown in the inset. Dark treatment results in the delivery of ATI1-labeled bodies carrying ER-marker to the vacuole (see inset in B). White arrowheads point to puncta where GFP and mCherry signals overlap. Chlorophyll autofluorescence is shown in magenta. Scale bars, 10μm. **(C)** Leaves of ATI1-GFP / mCherry-HDEL plants were either darkened or treated with 15 mg/ml TM for 24 h, and release of free GFP was monitored by analysis of total protein extracts with anti-GFP antibodies. Similar loading is shown by the stained level of ribulose bisphosphate carboxylase small subunit (loading). **(D)** Quantification of the ratio of free GFP to total GFP (free + fused to ATI1) demonstrate that ATI1 is degraded via autophagy in response to dark but not to TM treatment. Bars represent the mean ± SE (n=3); asterisk indicates a value significantly different from the control light sample; p < 0.01 (student’s t-test).

### ATI1 ER-phagy is TOR-dependent

The evolutionarily conserved protein kinase, Target of Rapamycin (TOR), integrates nutrient, stress and energy-related signals and is a main regulator of growth and cellular homeostasis. The TOR signaling pathway negatively regulates autophagy in yeast, mammals and plants [43]. Recently, TOR was shown to regulate autophagy induced by nutrient, salt or osmotic stress in Arabidopsis plants. However, autophagy induced by ER or oxidative stress was found not to be regulated by TOR signaling [44]. To analyze the role of TOR in the ATI1 autophagic pathway we utilized AZD8055, the strongest 2^nd^generation active-site TOR inhibitor in Arabidopsis [45, 46], and examined its effect on the accumulation of ATI-bodies. Seedlings stably expressing ATI1-GFP under its native promoter [38] were grown on solid ½ MS medium for a week and then transferred to liquid medium plus sucrose in the light, or minus sucrose in the dark in the presence of conA for 24 h. AZD8055 was added for the last 6 h of incubation. To reduce the number of ATI-bodies that accumulate under the dark treatment, a lower concentration of conA was used in these experiments compared to the experiments described in Figure 1. Representative images are shown in Figure 2A. Few ATI1-bodies were observed in seedlings growing in light plus sucrose, whether or not they were treated with AZD8055. However, treatment with AZD8055 significantly increased the number of ATI-bodies in dark treated seedlings (Figure 2B), suggesting that the ATI1 autophagic pathway is TOR-dependent.

**Figure 2.**
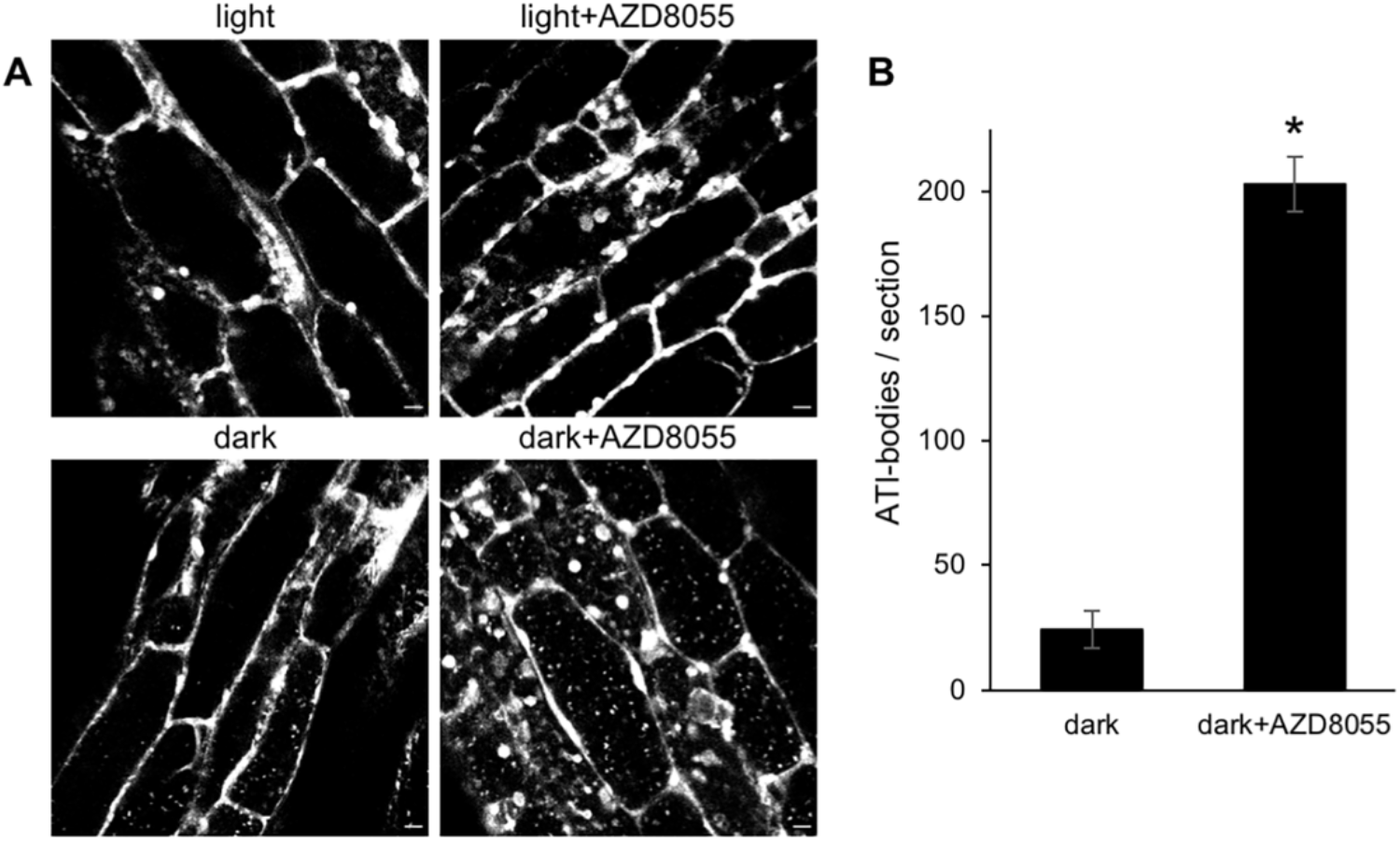
ATI1 autophagy is TOR-dependent. **(A)** Representative confocal images of *Arabidopsis* seedlings stably expressing ATI1-GFP (in green) under its native promoter that were grown on solid ½ MS medium for 7 days and then transferred for 24hr to liquid medium supplemented with sucrose in the light, or without sucrose in the dark in the presence of a low concentration of conA (0.5 mM) with or without addition of the TOR inhibitor AZD8055 for the last 6h. Chlorophyll autofluorescence is shown in magenta. **(B)** Quantification of ATI1-GFP labeled puncta in hypocotyl cells (section=0.04mm2) shows that addition of AZD8055 significantly induced the accumulation of ATI1-labeled puncta. Bars represent the mean ± SE (n=25 and 26 images from 7 plants for dark and dark+AZD8055 treatments, respectively); asterisk indicates a value significantly different from the control; p < 0.01 (student’s t-test). Scale bars, 10μm.

### The ATI proteins interact with MSBP1

To identify possible ER membrane-related protein interactors of the ATI proteins a split-ubiquitin yeast two-hybrid (Y2H) system was used. ATI2, as a representative, was fused to the C-terminal fragment of ubiquitin (Cub) along with an artificial transcription factor, and used as bait against a commercial library (pNubG-X) of Arabidopsis cDNA fused to the modified N-terminal fragment of ubiquitin (NubG). Several positive cDNA clones were detected in the Y2H screen, and were further analyzed by sequencing of the corresponding plasmids. Among the positive cDNA clones, Membrane Associated Progesterone Receptor 4 (MAPR4) appeared 5 times. One on one Y2H assay verified that MAPR4 interacted with both ATI1 and ATI2 (Figure 3A). The MAPR family is widespread in eukaryotes, and its members were shown to have varied cellular functions. *Arabidopsis* conatains four MAPR family members [47]. The best characterized is MSBP1 (MAPR5) that was shown to be involved in the signal transduction of the plant hormone brassinosteroid (BR) and in the redistribution of another major plant hormone, auxin. Moreover, MSBP1 was recently found to have a role in organizing and stabilizing cell-wall related lignin biosynthetic P450 enzymes on the ER [48–51]. Y2H assays demonstrated that similarly to MAPR4, MSBP1 also interacts with both ATI1 and ATI2 (Figure 3B). To confirm these results, *in vivo* BiFC assays were used [52]. Fusion proteins linking MSBP1 with the N-terminal fragment of the marker Enhanced Yellow Fluorescent Protein (EYFP; MSBP1-nEYFP), and ATI1 or ATI2 with the C-terminal fragment of the EYFP protein (ATI-cEYFP), were transiently co-expressed in *Nicotiana benthamiana* leaves. Interaction between the ATI proteins and MSBP1 is expected to bring the two halves of EYFP to close proximity and generate yellow fluorescence. Indeed, strong EYFP fluorescence resulted from the co-expression of MSBP1-nEYFP and ATI1 or ATI2-cEYFP, confirming the interaction between MSBP1 and the ATI proteins (Figure 3C). Similar results were obtained with MAPR4 (Figure S2).

**Figure 3.**
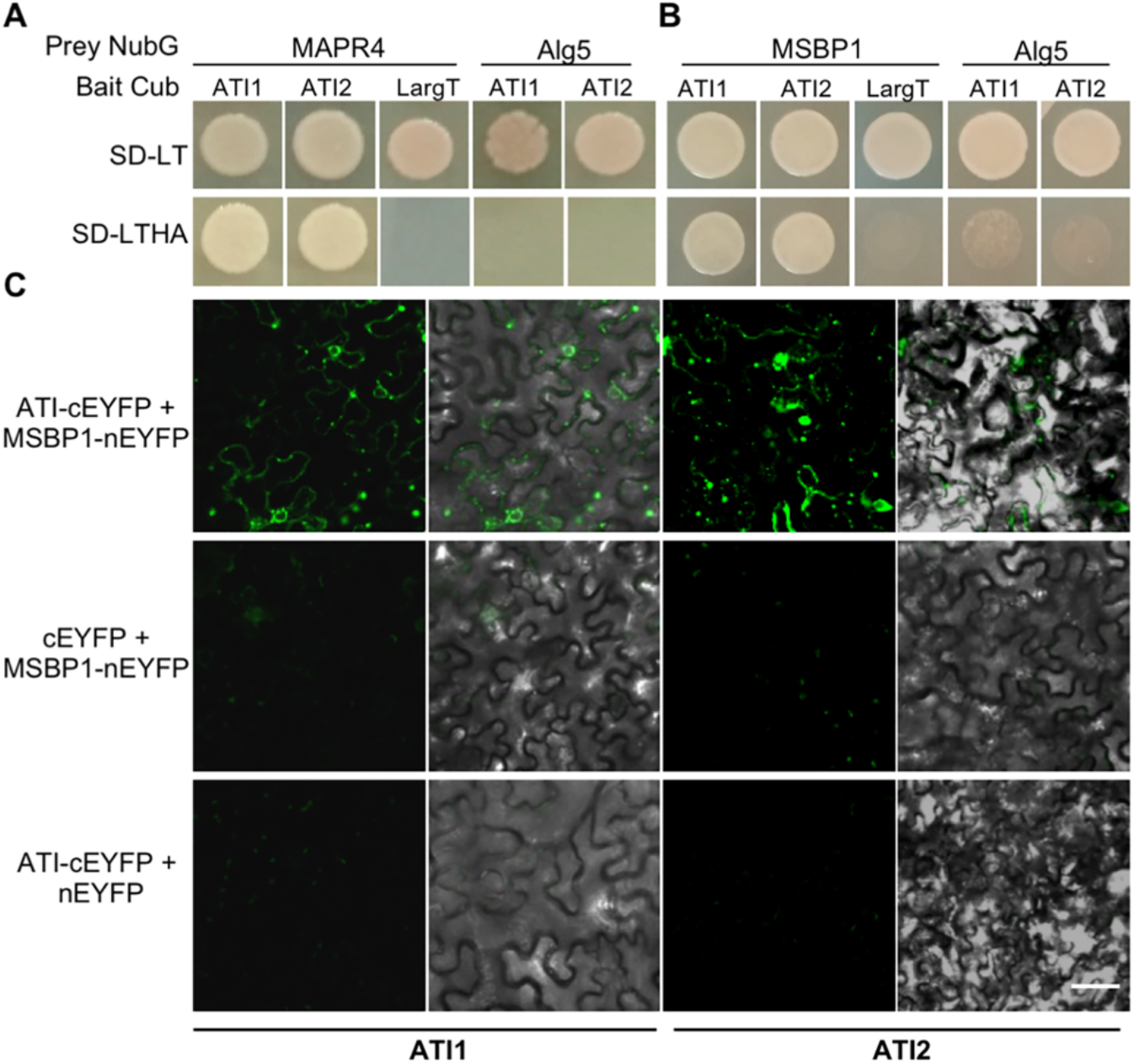
MAPR4 and MSBP1 interact with both ATI1 and ATI2. Split-ubiquitin yeast two-hybrid assay showing the interaction of MAPR4 **(A)** or MSBP1 **(B)** with ATI1 and ATI2. MAPR4 or MSBP1 were fused to the C-terminal fragment of ubiquitin (Cub) and ATI1 or ATI2 was fused to the modified N-terminal fragment of ubiquitin (NubG). The large T antigen (LargeT) and the endogenous ER protein Alg5 were used as negative controls. While all bait and prey couples can grow on non-selective media (SD-LT), the interaction between the ATI proteins and MAPR4 or MSBP1 brings the two halves of ubiquitin together and allows growth on selective media lacking histidine and adenine (SD-LTHA). **(C)** Confocal imaging of a BiFC assay involving co-expression of MSBP1-nEYFP with either ATI1 or ATI2 fused to cEYFP in *N. benthamiana* leaves (top panel). The yellow fluorescence signal indicates an interaction between the two proteins. The co-expression of ATI1 or ATI2-cEYFP with unfused nEYFP (middle panel) or the co-expression of MSBP1-nEYFP with unfused cEYFP (bottom panel) did not show any fluorescence. scale bars, 50μm.

### MSBP1 is degraded by autophagy

The interactions observed between MSBP1 and the ATI proteins suggest the involvement of autophagy in MSBP1 turnover. To further characterize the role of autophagy in MSBP1 degradation, we produced stable lines over-expressing mCherry-tagged MSBP1. Under regular growth conditions or in the absence of conA, very little mCherry signal was observed in the vacuoles of transgenic leaves. However, following dark incubation and treatment with conA, numerous MSBP1-mCherry labeled puncta resembling autophagic bodies were clearly visible in the vacuole (Figure 4A). To confirm that the MSBP1-mCherry labeled vacuolar puncta are autophagic bodies, we produced a line co-expressing MSBP1-mCherry with the GFP-ATG8f marker, that decorates autophagosomes from their formation to their lytic destruction in the vacuole [2, 53]. Upon dark and conA treatment, MSBP1-mCherry was indeed highly co-localized with GFP-ATG8f autophagic bodies in the vacuoles of both roots and leaves (Figures 4B and S3A). Furthermore, when MSBP1-mCherry was expressed in autophagy-deficient *atg5-1* mutant plants [54], no MSBP1-mCherry labeled puncta were observed under dark treatment and the addition of conA (Figure 4C). In agreement with the fluorescence microscopy results, the ratio of released free mCherry to MSBP1-mCherry substantially increased upon dark treatment in wild type plants, suggesting an increase in the autophagic flux. In contrast, very little free mCherry was observed in the *atg5-1* mutants, even following dark treatment (Figure 4D). Taken together, our results suggest that during dark-induced starvation, MSBP1 is delivered to the vacuole for degradation by autophagy.

**Figure 4.**
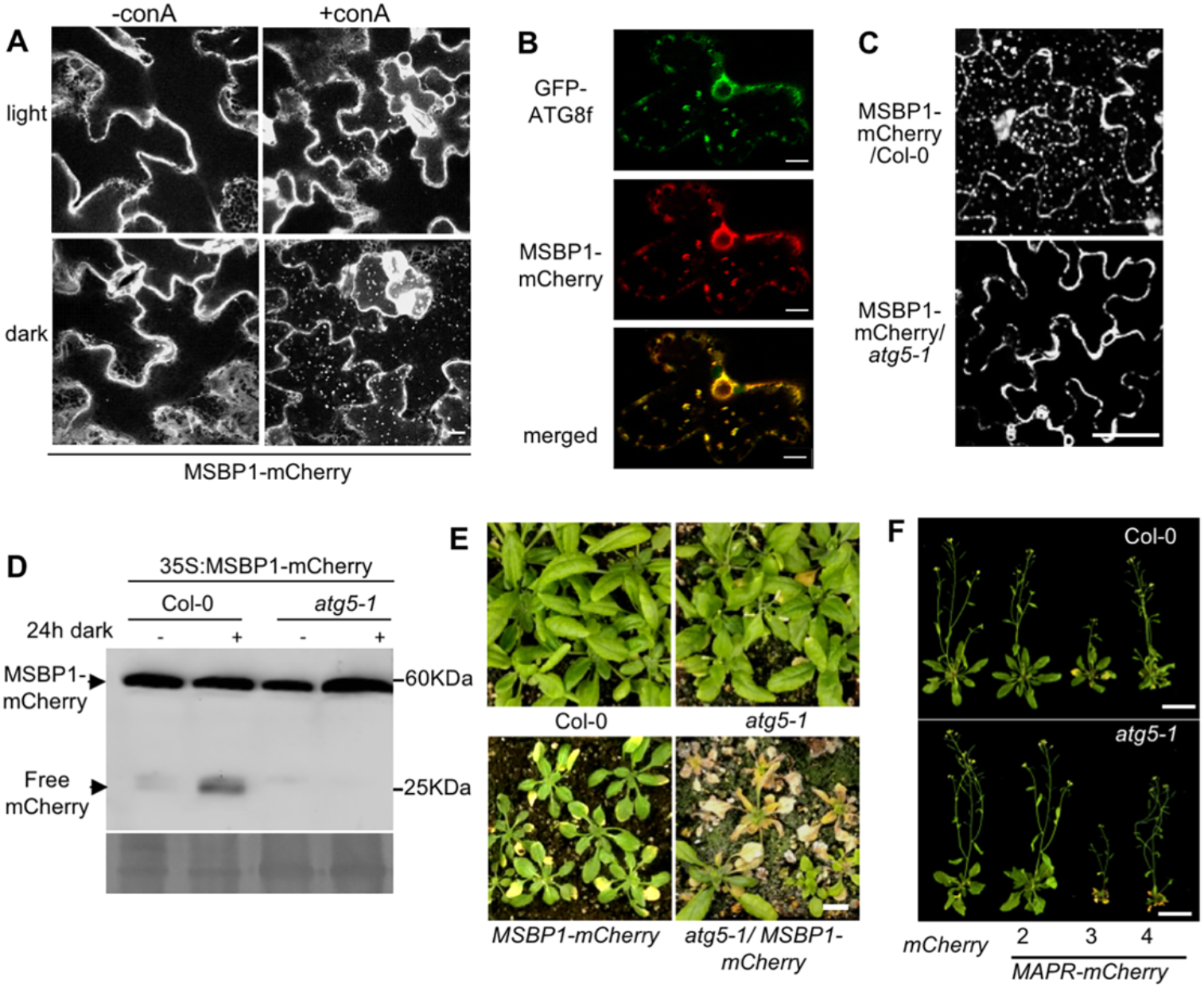
MSBP1 is associated with autophagic bodies and is degraded by autophagy. **(A)** Representative confocal images of leaves from plants stably expressing MSBP1-mCherry that were transferred for 16 h to liquid medium supplemented with sucrose in the light, or without sucrose in the dark with or without treatment with conA. MSBP1-mCherry labeled puncta accumulate in the vacuole following dark treatment in the presence of conA. Scale bar, 5μm. **(B)** Representative confocal images of roots of seedlings stably co-expressing MSBP1-mCherry and GFP-ATG8f following 16 h of dark and treatment with ConA. MSBP1-mCherry co-localizes with GFP-ATG8f in autophagic bodies. White arrowheads point to puncta where mCherry and GFP signals overlap. Scale bars, 5μm. **(C)** Representative confocal images of leaves from plants stably expressing MSBP1-mCherry in either Col-0 or *atg5-1* mutant backgrounds that were incubated for 16 h in the dark and treated with conA as in B. MSBP1-mCherry labeled autophagic bodies do not accumulate in the vacuoles of *atg5-1* mutant leaves. Scale bar, 25μm. **(D)** Leaves from plants stably expressing MSBP1-mCherry in either Col-0 or *atg5-1* mutant backgrounds were darkened for 16 h, and release of free mCherry was monitored by analysis of total protein extracts with anti-mCherry antibodies. The ratio of free mCherry to total mCherry (free + fused to MSBP1) increased in the Col-O, but not in *atg5-1* background, following dark treatment. Similar loading is shown by the stained level of ribulose bisphosphate carboxylase large subunit (RBCL) (loading). **(E)** Phenotypes of MSBP1-mCherry overexpressing lines in the background of Col-0 wild type plants or the *atg5-1* mutant. The phenotype of MSBP1 overexpression is exacerbated in the *atg5-1* background. Scale bars, 1cm. **(F)** Overexpression of MAPR3 (MSBP2) and MAPR4, but not MAPR2 shows early senescence phenotype. Scale bars, 2cm.

Interestingly the *atg5-1* autophagy deficient mutant background exacerbated the phenotype of MSBP1 over-expression. As previously observed, over-expression of MSBP1 in wild type plants led to reduced cell expansion resulting in smaller leaves [49, 51]. The over-expressing plants also showed early senescence phenotype (Figure 4E). Similar phenotypes were observed in plants over-expressing untagged MSBP1 (Figure S3B), and the severity of the phenotype was correlated with the level of over-expression of MSBP1 (Figure S3C). When MSBP1-mCherry was over-expressed in the *atg5-1* autophagy deficient mutant, the observed phenotype was much more severe, and most plants died before fruiting (Figure 4E). Among the four members of the Arabidopsis MAPR family, MSBP1, MAPR3 (MSBP2) and MAPR4, but not MAPR2, are predicted to have a transmembrane domain at the N-terminal of the protein [47]. Similarly to MSBP1, plants over-expressing mCherry-tagged MAPR3 (MSBP2) and MPAR4 had smaller leaves and showed early senescence phenotype, and the early senescence phenotype was aggravated in the background of the *atg5-1* mutant (Figure 4F). Similar phenotypes were not observed in plants over-expressing MAPR2, though the levels of the expressed proteins were similar in the different lines (Figures 4F and S4A), suggesting a different function for the non-membranal member of the family. Similarly to MSBP1, dark treatment induced accumulation of mCherry-labeled puncta in the leaf vacuoles of MAPR2, MAPR3 (MSBP2) and MAPR4-mCherry expressing plants, while no fluorescence was observed in the vacuoles of mCherry-MAPR expressing *atg5* mutants (Figure S4B). Hence, all MAPR family members, including the non-membranal MAPR2, are targeted by autophagy.

### MSBP1 is ER-localized and is delivered to the vacuole with an ER-marker

MSBP1 was initially reported to reside on plasma membrane and endocytic vesicles, and to negatively regulate BR signaling, affecting cell elongation and expansion [49–51]. However, a recent study demonstrated that MSBP1 as well as MAPR3 (MSBP2) are predominantly localized on the ER, where they function as scaffold proteins for lignin biosynthetic cytochrome P450 enzymes [48]. Indeed, when observed under confocal fluorescence microscope, mCherry labeled MSBP1 showed a reticulated pattern that co-localized with the ER marker GFP-HDEL, confirming its ER localization (Figure 5A left panels). Following carbon starvation and ConA application, MSBP1 also co-localized with the ER marker in autophagic bodies found within the vacuole lumen, suggesting that it is degraded by dark-induced ER-phagy (Figure 5A right panels).

**Figure 5.**
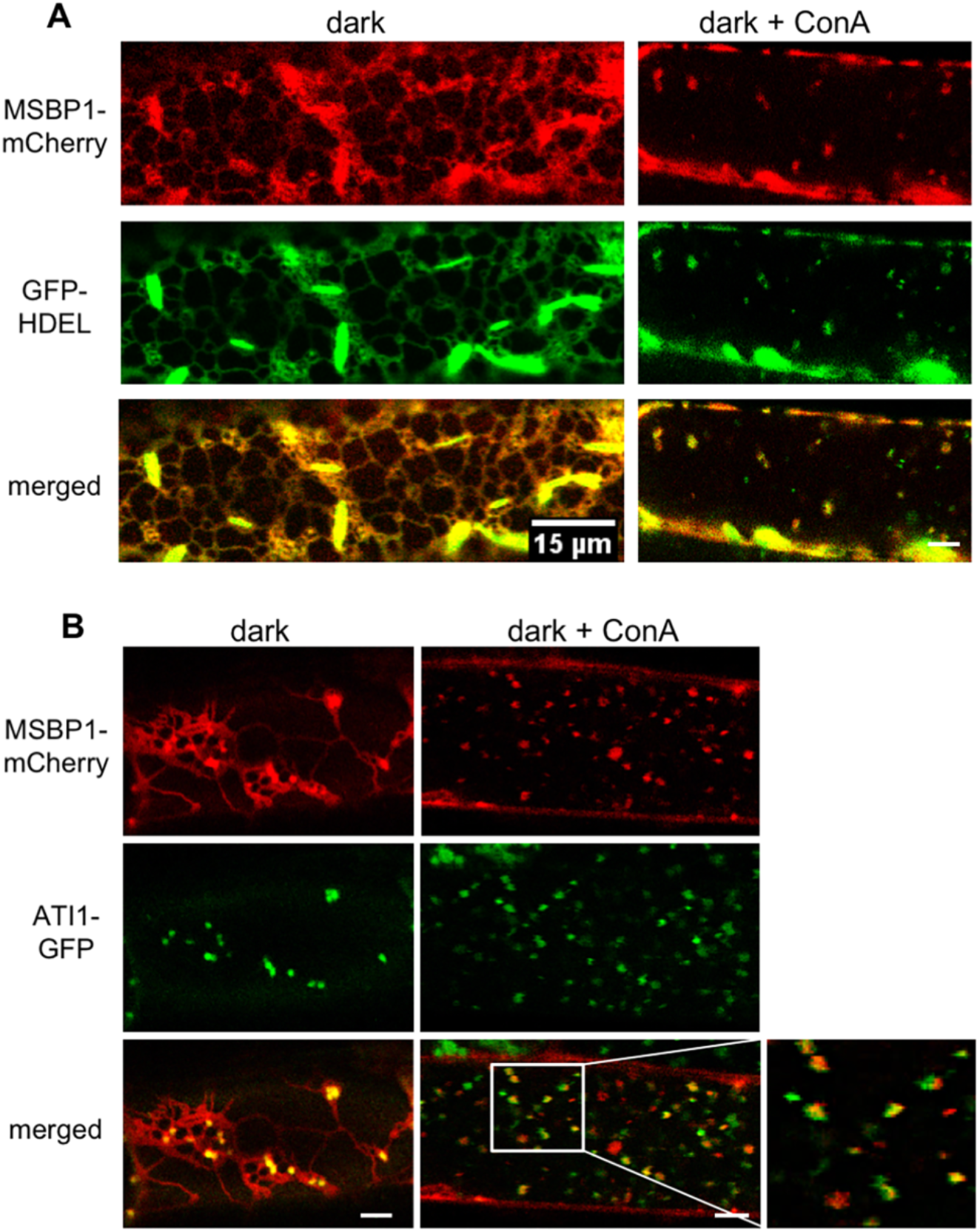
MSBP1-mCherry co-localizes with GFP-HDEL and ATI1-GFP in the ER and vacuoles. (A) Representative confocal images of root cells of seedlings stably co-expressing MSBP1-mCherry and GFP-HDEL show MSBP1-mCherry in red, GFP-HDEL in green and the overlay indicating co-localization in yellow (merged). 7-days old seedlings were transferred to liquid medium and incubated 16 h in the dark without sucrose. MSBP1-mcherry co-localizes with GFP-HDEL on the ER (left panel), and following treatment with ConA also in vacuolar puncta (right panel). **(B)** Similar treatment of seedlings stably co-expressing MSBP1-mcherry and ATI1-GFP show co-localization of MSBP1 and ATI1 on the ER (left panel) and on puncta in the vacuole (right panel). Magnification of the area in the white rectangle is shown on the right. Scale bars, 5μm.

### ATI1 is required for the autophagic degradation of MSBP1

As MSBP1 interacts with the ATI proteins (Figure 3) and both are targeted by ER-phagy in response to carbon starvation (Figure 1 and 5A), we wanted to look at the involvement of the ATI proteins in MSBP1 autophagic degradation. MSBP1-mCherry expressing lines were crossed with ATI1-GFP or ATI2-GFP expressing lines [37]. Seedlings co-expressing MSBP1-mCherry and ATI1- or ATI2-GFP were moved to growth medium without sucrose and kept in the dark for 24 h. Under these conditions, MSBP1-mCherry co-localized with both ATI1 and ATI2 labeled bodies in the cytosol (Figures 5B and S5, left panels). Following treatment with conA, MSBP1 co-localized with ATI1 and ATI2 in the vacuole as well (Figures 5B and S5, right panels). Furthermore, bodies co-labeled by ATI1 proteins and MSBP1 could be seen moving along the ER network and then entering the vacuole (Supplemental movie 1). To examine whether the ATI proteins are required for MSBP1 autophagic degradation, we produced a double knockout mutant of ATI1 and ATI2. CRISPR-Cas9 based technology was used to introduce an *ATI2* knockout mutation (as described in [40]) in the background of a homozygous SALK T-DNA insertion line (SALK_136644). MSBP1-mCherry was then expressed in the *ati1/2* mutant line. While numerous MSBP1-mCherry labeled autophagic bodies were visible in the vacuole of WT (Col-0) plants under carbon starvation and conA treatment, such bodies were significantly less visible in the ati1/2 background (Figure 6A). Quantification of MSBP-mCherry autophagic bodies corroborated the observed differences (Figure 6B). In accordance, analysis of free mCherry release compared to the full-length fusion protein in Col-0 compared to ati1/2 demonstrate a significantly reduced autophagic flux of MSBP1-mCherry in the background of three different ati1/2 lines (Figures 6C and D).

**Figure 6.**
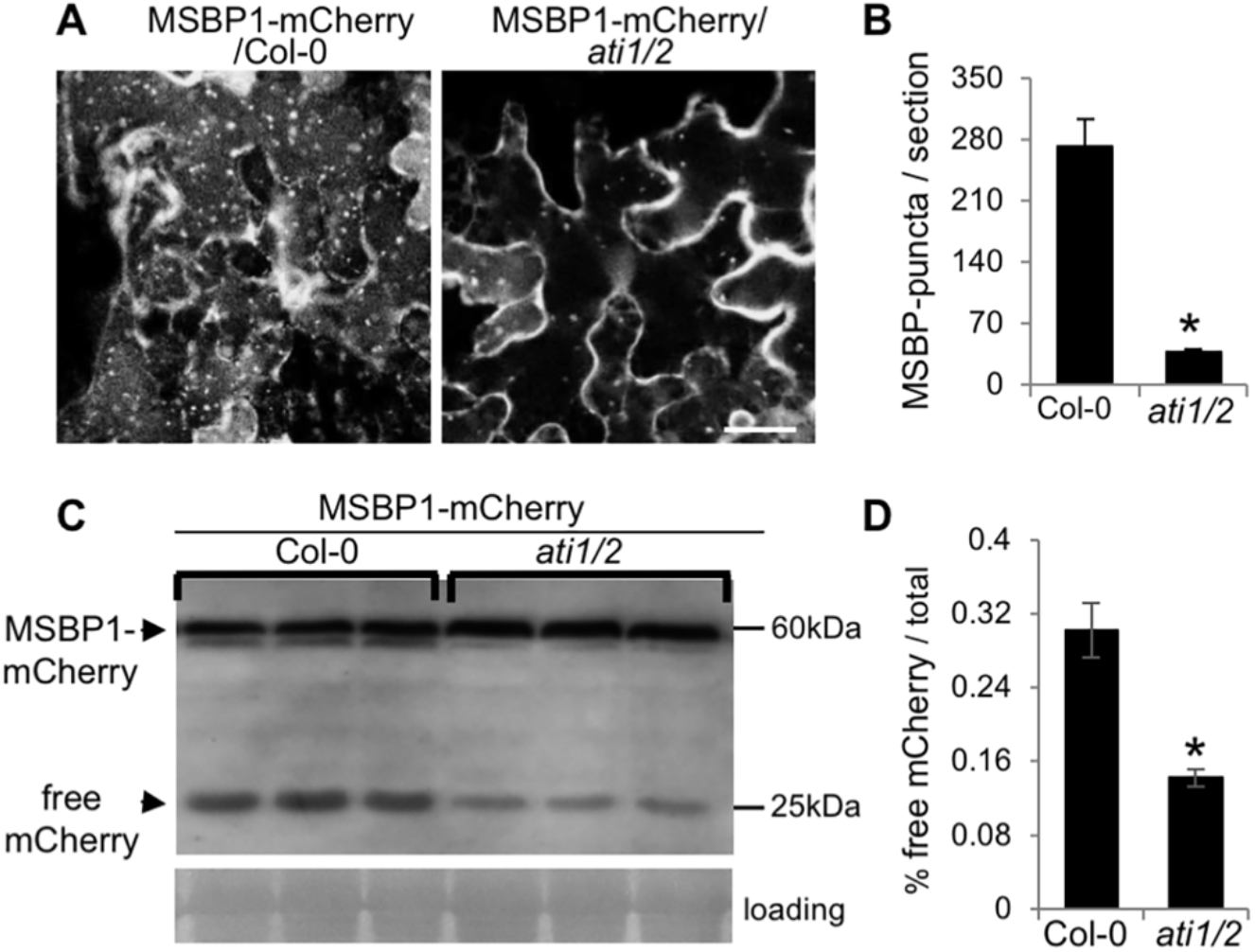
ATI is required for efficient augtophgic degradation of MSBP1. **(A)** Representative confocal images of leaves from plants stably expressing MSBP1-mCherry in the background of either Col-0 wild type (MSBP1-mCherry/Col-0) or ati1/2 double knockout plants (MSBP1-mCherry/ati1/2) that were incubated in the dark with conA for 16h. Scale bar, 20μm. **(B)** Quantification of MSBP1-mCherry-labeled puncta per section (0.005mm^2^) from Col-0 wild type or ati1/2 mutant plants. The number of dark-induced MSBP1-mCherry labeled puncta is reduced in ati1/2 knockout plants. Values represent mean ± SE (n=30 images from 5 different plants). Asterisk indicate statistically significant differences (P < 0.01, Student’s t-test). **(C)** Leaves of MSBP1-mCherry/Col-0 or MSBP1-mCherry/ati1/2 were dark treated, and release of free mCherry was monitored by analysis of total protein extracts with anti-mCherry antibodies. Similar loading is shown by the stained level of RBCL (loading). **(D)** Quantification of the ratio of free mCherry to total mCherry (free + fused to MSBP1) from the experiment shown in panel (C). Bars represent the mean ± SD. Asterisk indicate a statistically significant difference compared to Col-0 (*p*<0.01, Student’s t test).

## Discussion

The essential role of the ER in maintaining cellular homeostasis through the biosynthesis and transport of macromolecules, as well as its role in cellular signaling, requires constant adaptation of its size and activity to environmental and developmental cues. The identification and characterization of several mammalian ER-resident autophagy cargo receptors in the last few years suggests that selective ER-phagy is a major ER quality control mechanism under different abiotic and biotic stresses [3, 55]. However, our understanding of plant selective ER-phagy mechanisms is still limited. In this study we characterized a carbon starvation-induced selective ER-phagy pathway in *Arabidopsis* plants that involves the trans-membranal ATI proteins. The plant-specific ATI1 and ATI2 proteins were previously shown to localize to the ER under favorable growth conditions, and to associate with bodies that move along the ER and are transported to the vacuole upon carbon starvation in an autophagy-machinery dependent mechanism [37]. However, a direct link to ER-phagy was not demonstrated. Here we show that following dark treatment, the plant equivalent to starvation, ATI1-labeled bodies that carry an ER marker are transported to the vacuole, demonstrating the involvement of ATI1 in ER-phagy. Interestingly, ATI1 is not responsive to TM treatment, suggesting that the previously characterized ER-stress induced autophagy does not involve ATI1 (Figure 1) [24, 35, 56]. In agreement, and as generally shown for nutrient starvation induced autophagy in plants [44], ATI1 autophagy is TOR-dependent, as it is induced by TOR inhibition (Figure 2). As ER-stress induced autophagy was shown to be TOR-independent [44], this finding supports the conclusion that ER-stress and starvation-induced ER-phagy are mechanistically distinct. We further show that the ATI proteins interact with the ER-resident proteins MSBP1 and MAPR4 (Figures 3 and S2), and that these proteins, as well as other members of the MAPR family, are targeted by autophagy under carbon starvation (Figures 4, S3, S4). Interestingly, the autophagic degradation of MSBP1 is significantly reduced in ati1/2 mutant plants (Figure 6), suggesting that the ATI proteins function as selective ER-phagy cargo receptors to MSBP1.

The MAPR family of proteins is a found in yeast, mammals and plants and share a similar non-covalent heme/steroid-binding domain that is related to cytochrome b5. MAPR family members in mammals were shown to have a surprisingly diverse array of cellular functions including regulation of cytochrome P450, steroidogenesis, vesicle trafficking, progesterone signaling and mitotic spindle and cell cycle regulation [57]. Similarly, the best characterized MAPR family member in *Arabidopsis*, MSBP1, also seems to have multiple functions. MSBP1 was initially suggested to localize to the plasma membrane and to endocytic vesicles, and was shown to mediate the cycling of PIN2, a member of the PIN family of auxin efflux facilitators in the root [50], and to inhibit brassinosteroid (BR) signaling by enhancing the endocytosis of BAK1, a BR signal co-receptor [49, 51]. The inhibitory role of MSBP1 in BR signaling was also demonstrated in tobacco plants [58]. However, a recent paper suggested an additional localization and function for MSBP1 and MAPR3 (MSBP2). They were shown to reside predominantly on the ER, and to function as scaffold proteins to monolignol biosynthetic P450 enzymes, stabilizing their activity, thereby specifically controlling phenylpropanoid–monolignol branch biosynthesis [48, 51]. The ER localization of MSBP1 is supported by our results (Figure 5). Furthermore, we demonstrate that MSBP1 is specifically targeted by ER-pahgy as it is co-localized with an ER marker and with the ER-localized ATI proteins up to the vacuole (Figures 5 and S5). The identification of MSBP1 and the other MAPR family members as a target for autophagic degradation (Figures 4 and S4) is supported by a recent proteomics study showing that both MSBP1 and MAPR3 (MSBP2) are more abundant in *atg5* Arabidopsis mutants, though no change is seen in their transcription levels [59]. As MSBP1 was shown to serve as a physical scaffold to monolignol P450 monooxygenases, the whole complex might be directed to degradation through ATI-mediated ER-phagy under carbon starvation. Lignification requires a significant investment of fixed carbon and energy. Therefore, selective degradation of the lignin biosynthetic machinery under carbon starvation might be used to cope with energy deprivation. Finally, MSBP1 might be more than just a cargo of the ATI proteins, and might actually be involved in the regulation of ER-phagy. PGRMC1, a mammalian MAPR family member, was shown to bind LC3 and to promote autophagy [60]. In *Arabidopsis*, MAPR3 (MSBP2) was recently shown to interact with the ER stress regulator IRE1b and with GAAP1 and GAAP3, two Bax inhibitor-1 (BI-1)-like plant factors that play a role in plant survival under ER stress. Though the mechanistic details are not clear, these interactions were suggested to be involved in autophagy regulation in response to ER stress [61].

Similar to the known yeast and mammalian ER-phagy cargo receptors, the ATI proteins are ER membranal proteins that harbor an AIM or LIR (ATG8-Interacting Motif / LC3-Interacting Region) and can bind members of the ATG8 family [37, 55, 62, 63]. The known ER receptors were suggested to mediate selective autophagy of ER sub-regions based on their sub-ER localization [55]. Thus, the yeast Atg39 ER cargo receptor localizes to the perinuclear ER and induces autophagic sequestration of part of the nucleus, while yeast Atf40 is involved in the autophagy of the cytosolic and cortical ER [25]. In mammals, FAM134B was suggested to regulate the turnover of ER sheets, while RTN3L and ATL3 are important for degradation of ER tubules [27, 28, 31]. Though the sub-ER localization of the ATI protein is unknown, *Arabidospis* lignol p450 monooxigenases were shown to bind reticulons [64], a family of proteins that are needed to form the ER tubular network, most probably by stabilizing the high curvature of the tubules [65]. Hence, the association of the ATI proteins with MSBP1 might suggest that they are localized to ER tubules. Mammalian ER cargo receptors were suggested to function as a bridge between the ER and autophagosomal membranes, possibly through the Intrinsic Disordered Regions (IDRs) that are found in all of them [33]. Additionally, FAM134B and RTN3L might also function in reshaping or fragmenting the ER membranes in preparation for their engulfment by autophagosomes [27, 28]. The plant-specific ATI proteins also harbor an N-terminal IDR that contains their functional AIM motif [63]. However, ER-phagy involving the ATI proteins seems to have a distinct mechanism. Rather than direct association of cargo receptors on ER membranes with phagophores, this type of ER-phagy seems to involve first the formation of a distinct ATI-body, and only then fusion or engulfment by autophagosomes [37, 38]. As the ATI proteins are plant-specific, it will be interesting to see whether this mechanism is unique to plants. Moreover, while it is not clear how and if specific ER cargo is selected by the mammalian ER receptors, our results, together with very recent results from Michaeli et al. [40], suggest that the ATI proteins directly bind specific protein cargoes (MSBP1, MAPR4 and AGO1) and travel with them to the vacuole. Interestingly, in contrast to the autophagy mutant atg5, MSBP1 transport to the vacuole was significantly reduced, but not abolished, in the ati1/2 mutant (Figures 4C and 6). Thus, though the autophagy machinery is necessary for MSBP1 degradation, there seem to be some redundancy in the function of the cargo receptors. Quantitative proteomic analysis suggested partial functional redundancy between mammalian ER-phagy receptors [32]. It is likely that plant cells also have a repertoire of ER-phagy receptors, that are yet to be characterized, that together orchestrate ER-phagy under different stresses and in different ER sub-regions.

## Materials and Methods

### Plant materials and growth conditions

*Arabidopsis thaliana* Col-0 ecotype as well as *N. benthamiana* (for BiFC assays) were used in this study. The T-DNA insertion line *ati1* (SALKseq_136644) was obtained from the *Arabidopsis* Biological Research Center (ABRC) and its genotype was confirmed by PCR analysis. The *ati2* null mutant was generated by CRISPR-Cas9 editing [40], and the two lines were crossed to create the ati1/2 double mutant. The *Arabidopsis* mutant line *atg5-*1 (SAIL_129B079), and the transgenic lines GFP-HDEL, mCherry-HDEL, GFP-ATG8f, ATI1-GFP, ATI2-GFP, pATI1:ATI1-GFP and ATI1-GFP/mCherry-HDEL were previously described [37, 38, 40, 54, 66]. For MSBP1 expression, wild type plants or the appropriate lines were floral-dipped with *Agrobacterium tumefaciens* (strain GV3101) harboring 35S:MSBP1 or 35S:MSBP1-mCherry [67]. For seedlings experiments, seeds were surface sterilized using 3 % bleach, sown on 1/2 MS medium containing 1 % sucrose with the proper antibiotic selection, incubated in the dark at 4°C for 2-3 days and grown under long-day conditions (16 h light/8 h dark).

### Plasmid construction

To construct the plasmids used for the Yeast Two-Hybrid assays, BiFC assays and plant transformation, fragments containing the full-length CDS were digested and ligated into respective vectors. For Yeast Two-Hybrid assays, the pDHB1 and pPR3-N vectors were used for bait and prey constructs respectively. The different genes were ligated with the plasmids following SfiI digestion. For MSBP1 expression or BiFC plasmids construction, amplified DNA fragments were inserted into the appropriate pSAT vectors, carrying either full length mCherry or the N-terminal or C-terminal of EYFP [52, 68] using the In-Fusion kit (Clontech) according to the manual instructions. Modified pART27 or pBART binary vectors were produced by inserting into the NotI restriction site a multicloning site containing 13 unique restriction endonuclease recognition sites taken from the pPZP-RCS2 [52]. All generated pSAT expression cassettes were transferred to these modified pART27 or pBART binary vectors using either the AscI (for pSAT1) or the PI-PspI (for pSAT6) sites [52]. All the genes, plasmids, primers and the cleavage sites are listed in Supplementary Table S1.

### Split-ubiquitin yeast two-hybrid assays and BiFC

The Y2H screen was performed using the DUALhunter kit (Dualsystems Biotech) two-hybrid system. The ATI2 coding sequence was cloned into the yeast bait vector pDHB1 (Dualsystems Biotech), and co-introduced with a commercial pNubG-X *Arabidopsis* cDNA library (Dualsystems Biotech) to the NMY51 yeast strain. The NMY51 yeast strain contains the reporter genes ADE2, HIS3, and lacZ in its genome. The reporter genes are expressed only when the two-hybrid interaction combines the Cub and NubG parts of the yeast ubiquitin protein, resulting in the release of the LexA-VP16 transcription factor, which in turn enter to the nucleus and activates the three reporters. Positive colonies were selected on medium lacking the amino acids tryptophan, leucine, histidine, and aminodecanoic acid (SD - LTHA) supplemented with 10mM 3-aminotriazole (3-AT), a competitive inhibitor of the HIS3 gene product. The chosen clones were separately sequenced by using the forward library sequencing primer, 5’-GTCGAAAATTCAAGACAAGG-3’.

For analyzing in planta interactions between ATI1 or ATI2 and MSBP1 or MAPR4, different *Agrobacterium* strains harboring each of the following plasmids were used: pART27-ATI1-cEYFP, pART27-ATI2-cEYFP, pART27-MSBP1-nEYFP or pART27-MAPR4-cEYFP and the negative control plasmids pART27-nEYFP or pART27-cEYFP. The appropriate plasmid pairs were transiently co-transformed in *Nicotiana benthamiana* leaves following their infiltration, as previously described [69].

### Confocal Microscopy

A Nikon A1 confocal microscope system was used in this study. Generally, samples were put between two microscope glass cover slips (No.1 thickness) in an aqueous environment. For *N. benthamiana* transient expression, a small piece (0.5 cm2) of the injected leaf was placed between two glass cover slips as described above, and the epidermis cells were analyzed. For image acquisition, either the x20 or the x60 objectives (numerical apertures of 0.75 and 1.20, respectively) were used. GFP fluorescence images were taken using 488-nm laser excitation and the emission was collected via the 525-nm filter. The mCherry images were taken using 561-nm laser excitation and emission was collected through the 595-nm filter. YFP images were taken with the same settings as for GFP, and chlorophyll autofluorescence was imaged using the 640-nm laser and collected with the 700-nm filter. Time-lapse images were all composed from images taken using line sequence. Acquired images were analyzed using the NIS-elements AR imaging software. Quantification of ATI-bodies or MSBP1-mCherry labeled puncta was done manually.

### Treatments with chemicals

For ConA treatment, seedlings were grown in ½ MS agar Petri plates without sucrose for 6 d. Then, several whole seedlings of each of the examined lines were transferred into wells of 24-well plates containing ConA for diluted in liquid ½ MS medium. As a control, the same number of seedlings was subjected to the same treatment using DMSO (used to dissolve ConA). Seedlings were incubated in the liquid media at 23°C with gentle shaking (80 rpm). For experiments with ATI1-GFP/mCherry-HDEL, seedlings were incubated with 1mM conA for 24 h, while for experiments with the MSBP1 expressing lines, seedlings were incubated with 2mM conA overnight. Following incubation, the seedlings were directly examined by confocal microscopy. For leaf infiltration, newly matured rosette leaves were excised and infiltrated with 2μM ConA with a needleless syringe, and then wrapped with aluminum foil in a box paved with wet paper and incubated overnight at 23°C in the dark.

For dark and TM treatments of ATI-GFP/mCherry-HDEL mature plants, leaves of 4-5 week old plants were infiltrated with a needleless syringe with either 1μM ConA or with 15 mg/ml TM (stock solution of 10 mg/ml in DMSO). Leaves of control plants were infiltrated with the appropriate amount of DMSO alone. The plants were left for 24 h under regular light conditions or in the dark, as detailed in the figure legends, and then the treated leaves were either imaged by a confocal microscope or collected for protein extraction and immunoblotting.

For AZD8055 treatment, seedlings were grown as detailed above, and then incubated in liquid media supplemented with 0.5μM ConA for 24 h under regular light conditions or in the dark as detailed in the figure legend. AZD8055 (5μM) was added for the last 6 h, followed by confocal imaging.

### Protein extraction and western blot analysis

*Arabidopsis* leaves were ground to a fine powder in liquid nitrogen and proteins were extracted using extraction buffer (100 mM Tris-HCl pH 7.5, 1 % SDS, 5mM EDTA and 1× protease inhibitor cocktail). Following centrifugation at 12,000 rpm for 10 min at 4 °C, the supernatant was transferred to a new tube and quantified by BCA protein assay (Pierce). Reducing sample buffer was added, and the protein extract was separated on reducing SDS-PAGE and probed with commercial anti-RFP or anti-GFP antibodies (6G6, Chromotek or ab290, Abcam, respectively) to detect mCherry tagged MSBP1 or GFP-tagged ATI1. Band quantification was done using ImageJ.

### RNA analysis

Total RNA was extracted from newly matured leaves using a NucleoSpin RNA kit from MACHEREY-NAGEL according to the manufacturer’s instructions and then treated with DNase I. cDNAs were prepared from 2μg DNase-free total RNA using an oligo(dT)15 primer and a SuperScript II reverse transcriptase kit (Thermo Fisher Scientific) following the manufacturer’s instructions. Quantitative real time PCR was conducted using Fast SYBR Green Master Mix (Thermo Fisher Scientific) and the appropriate primers in an Applied Biosystems StepOnePlus Real-Time PCR thermocycler. *ACTIN2* was used as a reference gene. The primer sequences used for RT-qPCR can be found in Supplementary Table S1.

## Abbreviations

AGO1: ARGONAUTE1
ATI: ATG8-Interacting Protein
BiFC: Bimolecular Fluorescence Complementation
BR: brassinosteroid
conA: concanamycin A
DMSO: dimethyl sulfoxid
DTT: dithiothreitol
ER: endoplasmic reticulum
GFP: green fluorescent protein
MAPR: Membrane Associated Progesterone Receptor
MSBP: Membrane Steroid Binding Protein
SD: standard deviation
TM: tunicamycin
TOR: target of rapamycin
Y2H: yeast two hybrid

## Acknowledgments

The work was supported by a grant from the Israel Science Foundation (612/16). G.G. is the incumbent of the Bronfman Chair of Plant Sciences at The Weizmann Institute of Science.

