## Supplemental figures for "ATG8-interacting (ATI) 1 and 2 define a plant starvation-induced ER-phagy pathway and serve as MSBP1 (MAPR5) cargo-receptors"

### Supplemental Data:

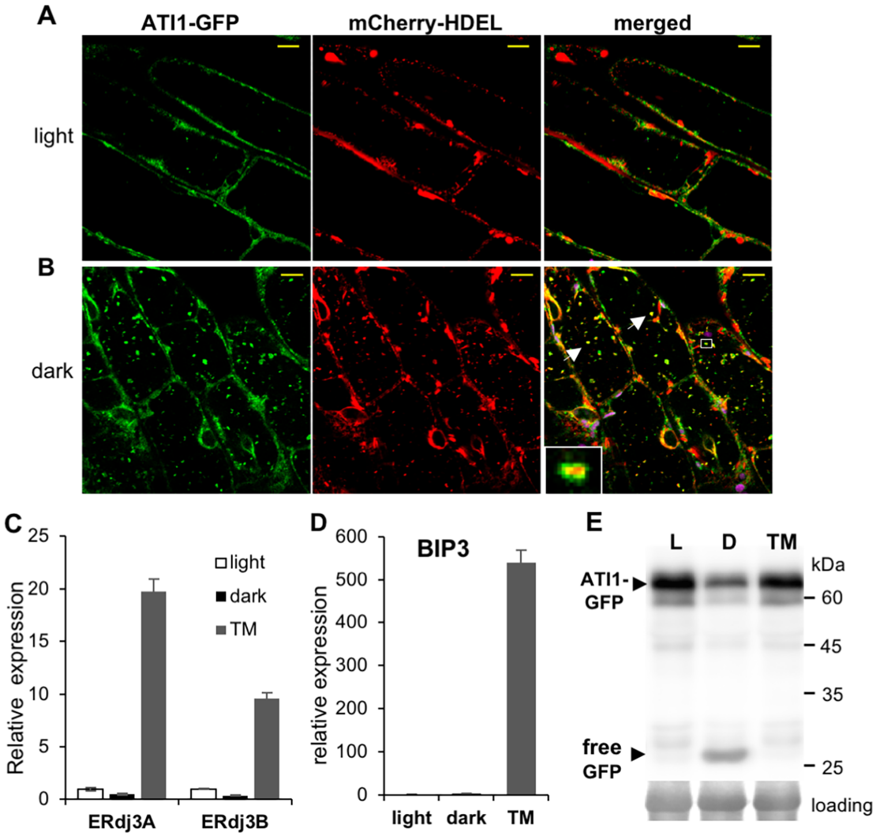


**Supplemental Figure 1. ATI1 is involved in ER-phagy in response to carbon starvation but not to ER stress.** Arabidopsis seedlings stably co-expressing ATI1-GFP and mCherry-HDEL were grown on solid ½ MS medium for 7 days and then either transferred to liquid medium supplemented with sucrose in the light **(A)** or to liquid medium without sucrose in the dark **(B)** for 24 h in the presence of conA. Representative confocal images of hypocotyl cells show ATI1-GFP in green, mCherry-HDEL in red and the overlay indicating co-localization in yellow (merged). Magnification of the area in the white rectangle is shown in the inset. Dark treatment results in the delivery of ATI1-labeled bodies carrying ER-marker to the vacuole (see inset in B). White arrowheads point to puncta where GFP and mCherry signals overlap. Scale bars, 10 μm. **(C and D)** ATI1-GFP/mCherry-HDEL co-expressing plants were either darkened, or left under regular light conditions and their leaves were infiltrated with 15 mg/ml for 24 h. The up-regulation of the well-established UPR target genes ERdj3A, ERdj3B **(C)** and Bip3 **(D)** was analyzed by quantitative RT-PCR. Data represent means ±SD for each condition (n=3). The obtained data were normalized against the expression of RNA Helicase (AT1G58050.1) and compared to the light condition. **(E)** Seedlings of ATI1-GFP / mCherry-HDEL plants were grown on solid ½ MS medium for 7 days and then either transferred to liquid medium supplemented with sucrose in the light with or without 15 mg/ml TM or to liquid medium without sucrose in the dark for 24 h. Release of free GFP was monitored by analysis of total protein extracts with anti-GFP antibodies. Similar loading is shown by the stained level of ribulose bisphosphate carboxylase small subunit (loading).

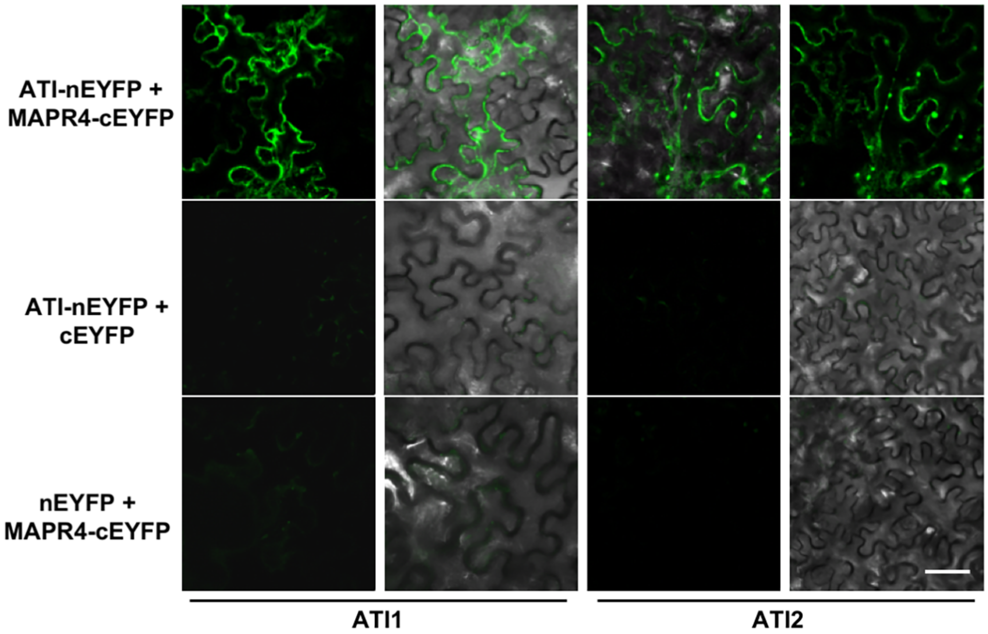


**Supplemental Figure 2. MAPR4 interacts with ATI1 and ATI2.** Confocal imaging of a BiFC assay involving co-expression of MAPR4-nEYFP with either ATI1 or ATI2 fused to cEYFP in *N. benthamiana* leaves (top panel). The yellow fluorescence signal indicates an interaction between the two proteins. The co-expression of ATI1 or ATI2-cEYFP with unfused nEYFP (middle panel) or the co-expression of MAPR4-nEYFP with unfused cEYFP (bottom panel) did not show any fluorescence. scale bars, 50 µm.

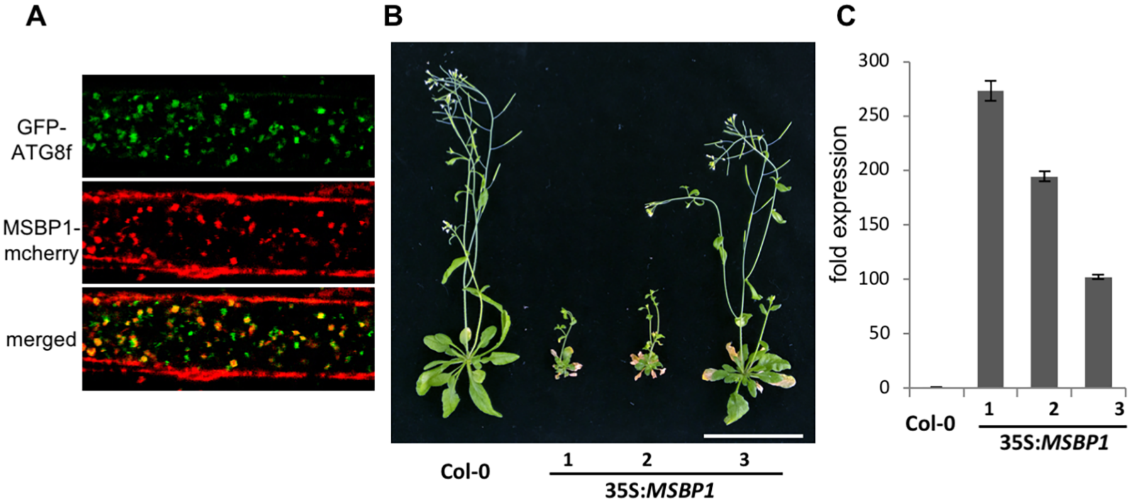


**Supplemental Figure 3.**

**(A)** Confocal images of conA treated leaves of plants co-expressing MSBP1-mCherry and GFP-ATG8f that were treated with conA and darkened for 16 h. MSBP1-mCherry co-localizes with GFP-ATG8f in autophagic bodies. Scale bar, 5µm. **(B)** Similarly to MSBP1-mCherry overexpressing plants, plants overexpressing untagged MSBP1 (*35S::MSBP1*) show early senescence phenotype. Scale bars, 5cm. **(C)** RT-PCR analysis of the relative expression of *MSBP1* in the transgenic lines shown in A. Phenotype severity is correlated with overexpression levels.

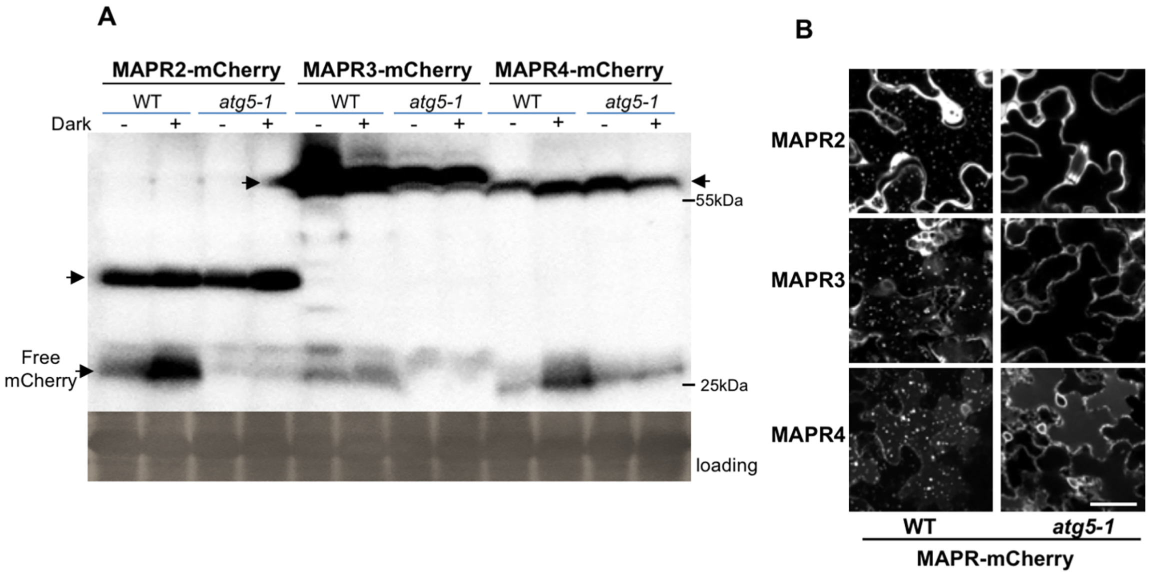


**Supplemental Figure 4. MAPR2, 3 and 4 are degraded by autophagy**
**(A)** Total protein extracts from plants stably expressing MAPR2, MAPR3 or MAPR4-mCherry in either Col-0 or *atg5-1* mutant backgrounds were analyzed with anti-mCherry antibodies. Similar loading is shown by the stained level of RBCL (loading). **(B)** Representative confocal images of leaves from plants stably expressing MAPR2, MAPR3 or MAPR4-mCherry in either Col-0 or *atg5-1* mutant backgrounds that were incubated for 16 h in the dark and treated with conA. Multiple mCherry labeled autophagic bodies accumulate in the vacuoles of Col-0, but not in the vacuoles of *atg5-1* mutant leaves. Scale bar, 25 μm.

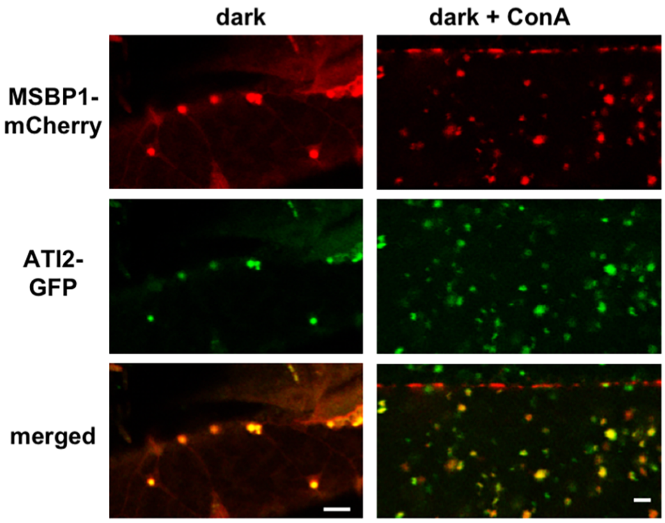


**Supplemental Figure 5. MSBP1 co-localizes with ATI2 in the ER and vacuoles.** Representative confocal images of root cells of seedlings stably co-expressing MSBP1-mCherry and ATI2-GFP show MSBP1-mCherry in red, ATI2-GFP in green and the overlay indicating co-localization in yellow (merged). 7-days old seedlings were transferred to liquid medium and incubated 16 h in the dark without sucrose. MSBP1-mcherry co-localizes with ATI2-GFP on the ER (left panel), and following treatment with conA also in vacuolar puncta (right panel). Scale bars, 5µm.
